# Express visuomotor responses reflect knowledge of both target location and conscious intent during reaches of different amplitudes

**DOI:** 10.1101/2022.11.06.515363

**Authors:** Samuele Contemori, Gerald E. Loeb, Brian D. Corneil, Guy Wallis, Timothy J. Carroll

**Author notes:** Submitting and corresponding author. The authors declare no competing financial interests.

## Abstract

When humans reach to visual targets, extremely rapid (~90 ms) bursts of activity can be observed on task-relevant proximal muscles. Such express visuomotor responses are inflexibly locked in time and space to the target and have been proposed to reflect rapid visuomotor transformations conveyed subcortically via the tecto-reticulo-spinal pathway. Previously, we showed that express visuomotor responses are sensitive to explicit cue-driven information about the target, suggesting that the express pathway can be modulated by cortical signals affording contextual pre-stimulus expectations. Here, we show that the express visuomotor system incorporates information about the veridical target-directed reaching metrics and contextual instructions during visuospatial tasks requiring different movement amplitudes. In one experiment, we recorded the activity from two shoulder muscles as participants reached toward targets that appeared at different distances. Longer hand-to-target distances led to larger and more prevalent express visuomotor responses than short-reach targets. This suggests that both the direction and distance of veridical hand-to-target reaches are encoded along the putative subcortical express pathway. In a second experiment, we modulated the movement amplitude by asking the participants to deliberately undershoot, overshoot, or stop (control) at the target. The overshoot and undershoot tasks impaired the generation of large and frequent express visuomotor responses, consistent with the inability of the express pathway to generate responses directed toward non-veridical targets (e.g. anti-reach tasks). Our findings appear to reflect strategic, cortically-driven modulation of the express visuomotor circuit to facilitate rapid and effective response initiation during target-directed actions.

**SIGNIFICANCE STATEMENT:** *Express* (~90 ms) arm muscle responses that are consistently tuned toward the location of visual stimuli suggest a subcortical contribution to target-directed visuomotor behaviour in humans, potentially via the tecto-reticulo-spinal pathway. This study shows that express muscle responses are modulated appropriately to reach targets at different distances, but generally suppressed when the task required non-veridical responses to overshoot/undershoot the real target. The data suggest that the tecto-reticulo-spinal pathway can be exploited strategically by the cerebral cortex to facilitate rapid initiation of effective responses during a visuospatial task.

## INTRODUCTION

Target-directed actions require knowledge of both the hand and target positions (Sabes, 2011; Proske and Gandevia, 2012). To catch a falling object for example, the multisensory information about the hand-to-target distance must be transformed into accurate motor signals to generate the muscle force and, in turn, accelerate the joints so that the hand can intercept the object before it hits the ground. Higher limb acceleration, therefore, results from greater activation of agonists and inhibition of antagonist muscles (Gordon et al., 1994).

Historically, target-directed visuomotor behaviour was thought to be the exclusive domain of cerebral cortex. This conception, however, is challenged by converging evidence showing that humans muscles can receive *express* target-driven motor signals at latencies (~90 ms) that leave little time for cortical visuomotor transformation (Goonetilleke et al., 2015; Gu et al., 2019; Selen et al., 2021; Billen 2022). Notably, the onset time of express visuomotor responses is far less variable than the mechanical reaction time (RT; Contemori et al., 2022), which depends mostly on longer latency (plausibly cortical) muscle response components. Express visuomotor responses are also inflexibly tuned to the physical stimulus location even in circumstances requiring volitional generation of non-veridical responses (e.g. anti-reach task; Gu et al., 2016). Critically, this suggests that this class of rapid target-directed responses is driven by the stimulus per se rather than a volitional command to start moving. This phenomenon, therefore, was proposed to reflect subcortical visuomotor transformations conveyed via the superior colliculus and its downstream reticulo-spinal circuit (Pruszynski et al., 2010).

Delineation of the factors that influence express visuomotor responses should provide clues about their origin and relationships to well-studied (putatively transcortical) visuomotor pathways. Previous work showed that the requirement to avoid rapid target-directed responses impaired the generation of express visuomotor responses (Pruszynski et al., 2010; Wood et al., 2015; Atsma et al., 2018). More recent work showed that express visuomotor responses are modulated by explicit cues about the temporal (Contemori et al., 2021a) and spatial (Contemori et al., 2021b) presentation of visual stimuli, and that express responses can incorporate contextual pre-stimulus expectations about the required movement to reach the target (Gu et al., 2018; Contemori et al., 2022). In all, these findings appear to reflect context-dependent modulation of the express circuit, potentially conveyed top-down via direct cortico-tectal (see for review Boehnke and Munoz, 2008) and cortico-reticular projections (Keizer and Kuypers, 1984, 1989; Fregosi et al., 2017; Darling et al., 2018; Fisher at al., 2021). Here we asked if express visuomotor responses are modulated compatibly with the required movement amplitude to accomplish a visuospatial task. If so, it would suggest that the circuits responsible for express limb activity produce control signals that account for the details of reach metrics, rather than merely the initial reach direction.

We conducted two experiments to explore express visuomotor responses to targets that required different movement amplitudes via modulation of: (i) the physical hand-to-target distance; (ii) the explicit instruction to overshoot, undershoot, or stop at the target. The first experiment showed that express visuomotor responses were facilitated by increasing the hand-to-target distance, suggesting that the express system encodes both the direction and distance metrics of veridical target-directed reaches. The second experiment showed significantly fewer and smaller express visuomotor responses, and longer volitional RTs, for both overshooting and undershooting tasks compared to veridical target-directed reaching actions. This suggests that express visuomotor behaviour is generally inhibited in circumstances requiring sensory-to-motor transformation for abstract targets; a task that is incompatible with the stimulus-locked output of the putative subcortical express circuit (Gu et al., 2016). The findings support the idea that the cerebral cortex strategically exploits the express pathway when its motor output is functional for rapid initiation of veridical target-directed actions, but suppresses the express network when it is incapable of meeting the current task demands.

## MATERIALS AND METHODS

### Participants

Fourteen adults completed the first experiment (6 females; mean age: 30.9±9 years), and twelve of them also participated in the second experiment (4 females; mean age: 31.8±9.2years). All participants were right-handed, had normal or corrected-to-normal vision, and reported no current neurological or musculoskeletal disorders. They provided informed consent and were free to withdraw from the experiment at any time. All procedures were approved by the University of Queensland Medical Research Ethics Committee (Brisbane, Australia) and conformed to the Declaration of Helsinki.

### Experimental set-up and task design

#### Experimental set-up

For both experiments, the participants performed visually guided target-directed reaches using a two-dimensional planar robotic manipulandum (the vBOT, Figure 1A; Howard and Ingram, 2009). In the vBOT setup, the visual feedback is provided via an LCD computer monitor (120Hz refresh rate) mounted above the robot handle and projected to the participant via a mirror, which occludes direct vision of the arm (Figure 1A). The visual stimuli were created in Microsoft Visual C++ (Version 14.0, Microsoft Visual Studio 2005) using the Graphic toolbox. The hand position was virtually represented by a blue cursor (~1 cm in diameter) whose apparent position coincided with actual hand position in the plane of the limb. The kinematic data of the vBOT handle were sampled at 1 kHz. The target was a filled black circle of 3 cm in diameter presented against a light grey background to create a high contrast stimulus (target luminance ~0.5 cd/m^2^, background luminance ~150 cd/m^2^; Cambridge Research System ColorCAL MKII). During the experiments, the upper arm was supported on a custom-built air sled positioned under the right elbow to minimize sliding friction (Figure 1A).

**Figure 1:**
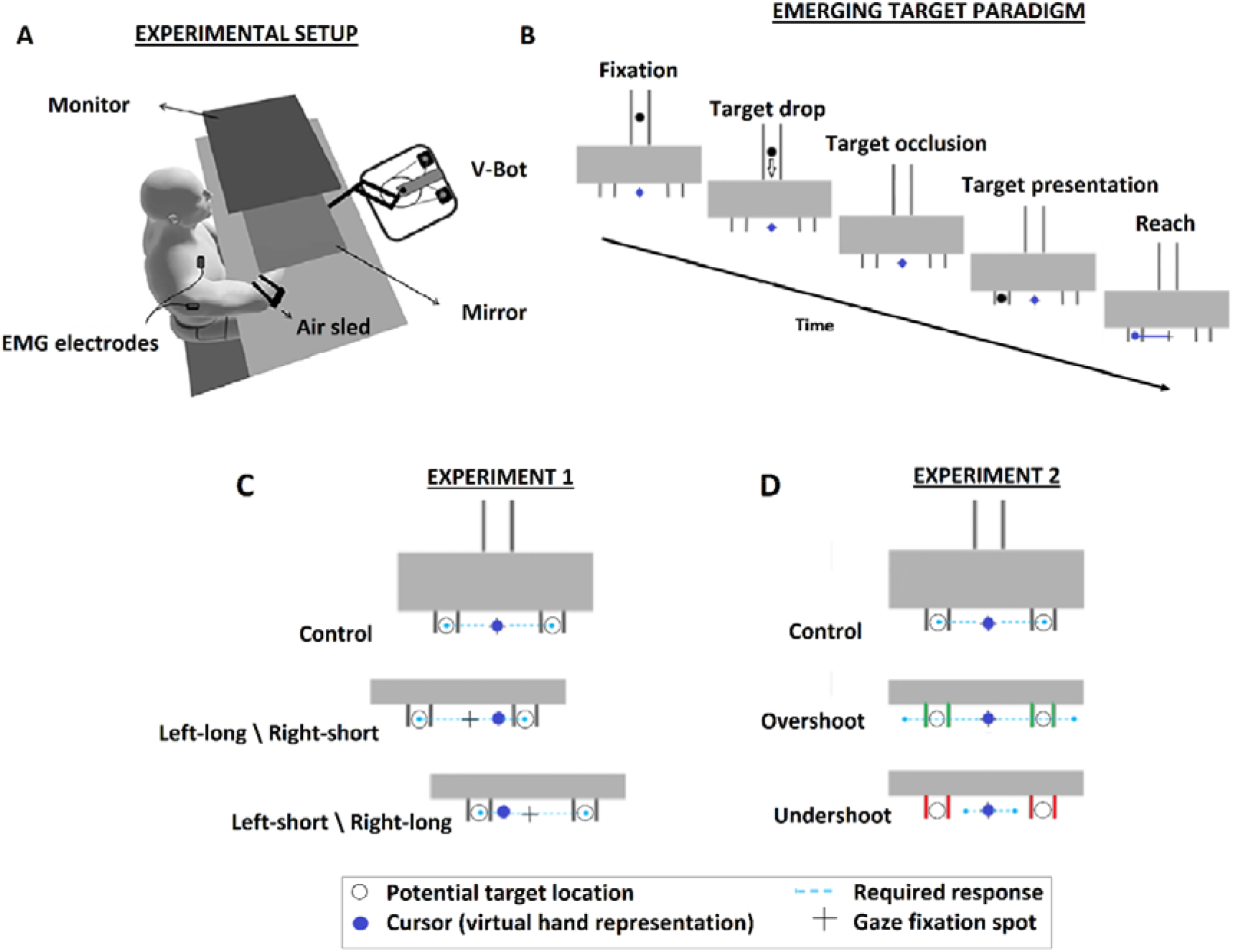
A: experimental setup. Participants’ hand positions were virtually represented via a cursor (blue dot in panels B-D) displayed on the monitor and projected into the(horizontal) plane of hand motion via a mirror. The head position was stabilized by a forehead rest (not shown here). B: schematic diagram of the timeline of events in the *emerging target* paradigm. Once the cursor was at the home position, the “+” sign for fixation was presented underneath the barrier. After one second of fixation, the target started dropping from the stem of the track at constant velocity of ~30 cm/s until it passed behind the barrier (i.e., occlusion epoch) for ~480 ms, and reappeared underneath it at ~640 ms from its movement onset time. C: task conditions in experiment one. In the control condition, the right and left potential target locations (unfilled grey circles underneath the barrier) had equal distance from the reaching hand. In the long-reach condition the target required a longer reach relative to control condition, whereas in the short-reach condition the target was closer to the reaching hand relative to control conditions. In these examples, shifting the visual elements toward the left required long and short reaching distances to address the location of left and right targets, respectively. By contrast, rightward shifts of the visual element generated the opposite direction-x-distance conditions of reaching. D: task conditions in experiment two. Here, the right and left target potential target locations (unfilled grey circles underneath the barrier) had equal distance from the reaching hand across all conditions. In the control condition, the vertical lines underneath the barrier were coloured black and the hand had to stop at the actual target location. By contrast, the hand had to overshoot or undershoot the actual target location when the vertical lines underneath the barrier were green (i.e. overshooting condition) or red (i.e. undershooting condition), respectively.

In both experiments, the target was presented via an *emerging moving* target paradigm (Figure 1B; Kozak et al., 2020; Kearsley et al., 2022; Contemori et al., 2022). To start the trial the participants had to align the cursor and gaze at a ‘home’ position (a blue ring of ~2 cm in diameter) located at the centre of the monitor and aligned with the mid-body line. Once the ‘hand-at-home’ condition was fulfilled, the ring was turned off and the fixation spot changed to a “+” sign. Note that for the first experiment the fixation spot position was not always coincident with the starting hand position, but rather changed as function of the trial condition (for details see Experiment 1: task design, and Figure 1C). Simultaneously with the fixation spot, a constant rightward load of ~5N was applied to preload the shoulder transverse flexor muscles, including the clavicular head of the pectoralis major muscle. At the same time, we also displayed the target close to the top of the monitor and within a track, shaped as an inverted diapason, whose rightward\leftward deviation point was occluded by a rectangular black barrier (Figure 1B). After ~1s of fixation, the target fell toward the bottom of the screen, disappeared behind the barrier and reappeared underneath it by making one single flash of ~8 ms of duration at the right or left of participants’ right hand and fixation spot. Notably, this paradigm allowed us to generate a high-contrast, low spatial-frequency, transient and temporally predictable stimuli that are all critical attributes to facilitate express visuomotor responses (Wood et al., 2015; Kozak et al., 2019; Contemori et al., 2021a). The participants were instructed to not break fixation until the target emerged from behind the barrier and to start moving as rapidly as possible toward the target. “Fixation” or “Too fast” errors were shown if the participants did not respect the gaze fixation requirements or moved before the target presentation, respectively, and the trial was reset.

Horizontal gaze-on-fixation was checked on-line with bitemporal, direct current electrooculography (EOG) sampled at 1 kHz. The time at which the stimulus was displayed on the monitor was recorded with a photodiode that detected a secondary light appearing at the bottom-left corner of the monitor and simultaneously with the actual target. Note that the photodiode fully occluded the secondary light thus making it invisible for the participants.

#### Experiment 1: task design

In the first experiment, we investigated modulations of express visuomotor responses as a function of the physical hand-to-target distance. To this aim, the participants had to reach a visual target whose distance from the hand was varied trial-by-trial to create three trial conditions: (i) *control-reach* condition, when the hand-to-target distance (~8 cm) was equivalent for both right and left targets; (ii) *long-reach* condition, when the hand-to-target distance was longer (~13 cm) than control; (iii) *short-reach* condition, when the hand-to-target distance was shorter (~3 cm) than control. The hand-to-target distance was modulated by shifting the target, track and visual barrier ~5 cm rightward, or leftward, relative to the static home position of the hand, such that separate long and short reaches were required for left and right targets (e.g. leftward shift → left-long/right-short reaches; Figure 1C). Note that the between-target distance (~16 cm) was kept constant, and the fixation point was shifted by ~4 cm such that the target induced the same retinal error across conditions. Also, the subjects always had >1s to perceive the shift of the visual elements to ensure unambiguous interpretation of the trial context before the target presentation.

To control the oculomotor behaviour, the EOG was calibrated before the main experiment by asking the participants to look at a target located at the centre of the monitor (consistent with the fixation spot location in control conditions; Figure 1C) for ~10s. Then the target jumped laterally right/left at three different distances (i.e. six direction-x-distance conditions), stayed there for ~2s before returning back to the initial one and made another jump only after another ~5s. For consistency with the main experiment, the target was a filled black circle 3cm in diameter presented against a light grey background and jumped ±8, ±13 and ±3 cm relative to the starting central position. The target jumped laterally five times for every direction and distance condition (i.e. 30 total trials). Importantly, this procedure allowed us to define the within-subject absolute EOG signal values across different eye positions and, thereby control the gaze fixation online.

For the main experiment, each participant completed 6 blocks of 48 reaches/block (24 for each direction), with each block consisting of 16 control-reach, 16 long-reach and 16 short-reach trials, randomly intermingled.

#### Experiment 2: task design

The first experiment showed modulations of express visuomotor response as a function of the reaching distance, suggesting express encoding of the physical target distance from the hand (for details see Experiment 1 results). Alternatively, the data might reflect context-based preparation of long\short movements independently of the real hand-to-target distance. Although these alternatives are not mutually exclusive, we ran a second experiment to test express visuomotor response in contexts requiring different target-directed movement amplitudes because of explicit instruction to: (i) stop at the target (*control*); (ii) *overshoot* the target; (iii) *undershoot* the target (Figure 1D). The control condition replicated that of the first experiment as the participants had to stop at the actual target location, or at least within the two vertical black lines where the target appeared underneath the barrier (Figure 1D: control condition). For the overshoot condition, we displayed green vertical lines underneath the barrier and instructed the participants to overshoot the actual target location by ending the movement at least beyond the outermost vertical green line (Figure 1D: overshoot condition). For the undershoot condition, we used red lines beneath the barrier and asked the participants to undershoot the actual target location by ending the movement before the innermost vertical red line (Figure 1D: undershoot condition). On every trial, the target always appeared at ~8 cm to the right or left of participants’ right hand. Note that the second experiment design did not require distinct movement amplitudes for different target locations (e.g. right-overshoot vs left-undershoot). The motivation for providing advance notice to stop, overshoot or undershoot both the right and left targets was to dissociate the executed reach from the target location without adding complexity for the trajectory-endpoint decision at the time of target presentation. To this aim, and consistent with the first experiment, the trial condition (i.e. the colour of the lines underneath the barrier) was made explicit to the participants for >1s before the target presentation.

Each participant completed 6 blocks of 48 reaches/block (24 for each direction), with each block consisting of 16 control-reach, 16 overshoot-reach and 16 undershoot-reach trials, randomly intermingled.

### Data recording and analysis

#### Kinematic data analysis

Consistently with our previous work (Contemori et al., 2022), we defined the mechanical reaction time by indexing the first time point at which the radial hand velocity exceeded the baseline mean velocity (i.e. average velocity recorded in the 100 ms preceding the target onset time) by more than five standard deviations (see Figure 2B for exemplar results). Trials with RT <140 ms (~3%) or >500 ms (<1%) were excluded from offline analysis (Contemori et al., 2021a, 2021b, 2022).

**Figure 2:**
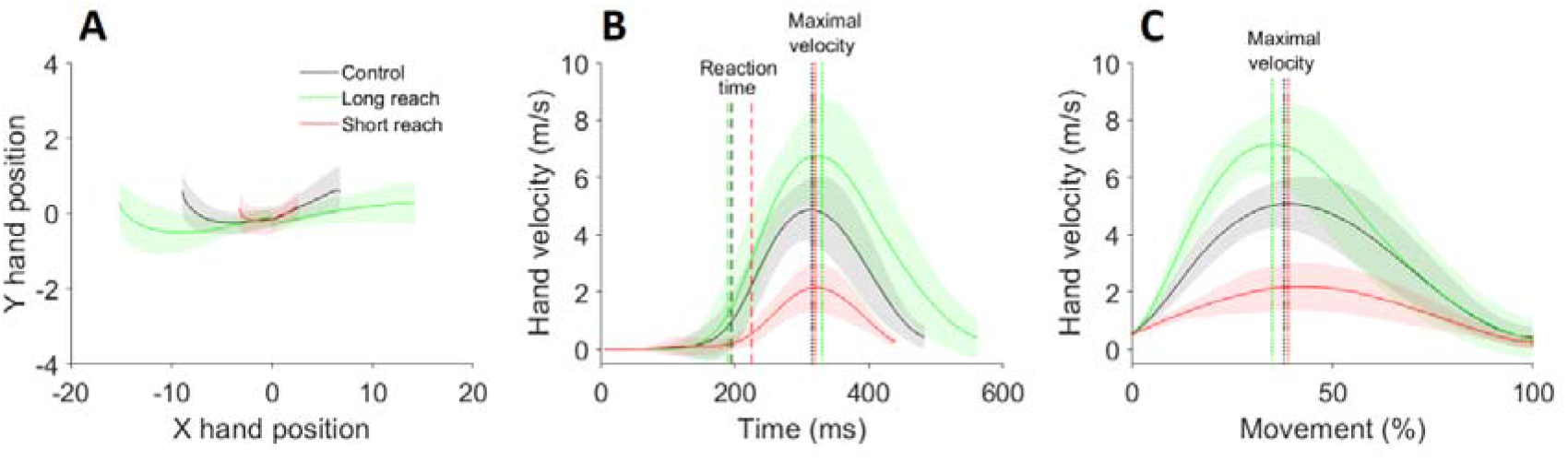
Kinematic data of an exemplar participants for the first experiment. A, hand trajectories in the control (black traces), long-reach (green traces) and short-reach (red traces) conditions. B, condition-dependent hand velocity traces. The time is relative to the target presentation. Vertical dashed and dotted lines are used to display the mechanical reaction times and maximal velocities across conditions, respectively. C, time-normalization of the hand velocity traces for the entire movement duration and point of the movement at which the peak velocity was reached. The data are plotted as mean (solid lines) and standard deviation (shaded area around the mean lines).

To determine the response correctness, we measured the initial reach direction by adopting a procedure previously described by Contemori et al., 2022. Briefly, we compared the initial hand-trajectory direction (i.e. slope of a line connecting the hand position coordinates at the RT and the 75% of the peak velocity) with the actual target location. We then computed the movement endpoint by searching for the point in time at which the total hand velocity fell below 0.5 m/s after having reached its peak value. We reasoned that a trial was correct if the hand initially moved toward the actual target and ended at the location specified by the trial-condition. We also computed the movement time (i.e. time difference between the movement-end and RT), and the time to reach the maximal velocity. To test whether the movement evolved similarly across conditions beyond differences in movement time, we conducted a trial-by-trial temporal normalization for the whole movement duration. This also allowed us to index the point within the movement at which the hand-velocity reached its peak. For both experiments, the kinematic data were averaged across the left and right directions to limit biases related to the preload.

#### EMG data analysis

Surface EMG activity was recorded from the clavicular head of the right pectoralis muscle (PMch) and the posterior head of the right deltoid muscle (PD) with double-differential surface electrodes (Delsys Inc. Bagnoli-8 system, Boston, MA, USA). The quality of the EMG signal was checked offline with an oscilloscope by asking the participants to flex (PMch activation-PD inhibition) and extend (PMch inhibition-PD activation) the shoulder in the transverse plane. The sEMG signals were amplified by 1,000, filtered with a 20–450 Hz bandwidth filter by the “Delsys Bagnoli-8 Main Amplifier Unit” and sampled at 2 kHz using a 16-bit analog-digital converter (USB-6343-BNC DAQ device, National Instruments, Austin, TX). Trial-by-trial, the quality of the EMG signal was verified online on the experimenter’s computer via a custom Matlab script that generated live plots of the recorded data. The sEMG data were then down-sampled to 1 kHz and full-wave rectified offline.

To detect the earliest stimulus-driven muscle response, we adopted a single-trial analysis method named the *detrended-integrated* signal method that we recently developed and validated (Contemori et al., 2022). Briefly, we computed the integral of the EMG trace for each millisecond recorded between 100 ms before and 300 ms after the target onset time. We then detrended the signal by subtracting the linear regression function of the background period (from 100 ms before to 70 ms after the stimulus presentation) from the entire 400 ms window and computed the background average and standard deviation values of the detrended-integrated signal. We indexed the “candidate” muscle response onset time as the first time the detrended-integrated signal exceeded the background average value by more (i.e. signature of muscle activation), or less (i.e. signature of muscle inhibition), than five standard deviations. Finally, we ran a linear regression analysis around the candidate muscle response onset time and searched for the interception point between the linear trendline the background value of the detrended-integrated signal. Notably, this allowed us to find the time the detrended-integrated signal started diverging from background. Consistent with previous work, we defined muscle responses in each trial as “express” if they initiated within 70-120 ms after the target presentation (Gu et al., 2016; Contemori et al., 2021a, 2021b, 2022).

One of the most distinctive attributes of the express visuomotor response is that its onset time does not reflect the trial-by-trial variability of long-latency responses because it is invariant at ~100 ms from stimulus presentation regardless of the mechanical RT (Pruszynki et al., 2010; Wood et al., 2015; Kozak et al., 2019, 2020; Kozak and Corneil, 2021; Contemori et al., 2021a, 2021b, 2022). Critically, the broad range of delays for a motor signal to reach the RT detection threshold is consistent with poly-synaptic nature of cortical sensorimotor networks to transform sensory inputs into deliberate decisions for actions. By contrast, the strikingly short-latency and relative temporal consistency of express visuomotor responses implies a small range of delays in motor signal conduction time, consistent with the few synapses of the tecto-reticulo-spinal pathway. To test the extent to which the express visuomotor response onset times were independent from the manual RT, we adopted a procedure previously described by Contemori et al., (2022) in which we first binned the express visuomotor response trials in “express-fast” and “express-slow” trial sets according to the median RT value of the full class of express trials. We then computed the average express response initiation time of the express-fast and express-slow trial sets as well as the average RT of the corresponding fast and slow trial sets. Finally, we fitted a line to the data to test if the muscle response onset time did not co-vary with the RT (i.e. line slope >67.5 deg; for details see Contemori et al., 2022; see also Figure 3 in Contemori et al., 2021a and 2021b). In this case, we computed the average response initiation time and prevalence (%) across the express visuomotor response trials. In this circumstance, we also quantified the express response magnitude by computing the average EMG activity recorded in the 10ms after the response initiation time for each rightward and leftward trial exhibiting an express visuomotor response. We then averaged this metric across the express response trials and computed the difference between the left and right targets (Contemori et al., 2022).

**Figure 3:**
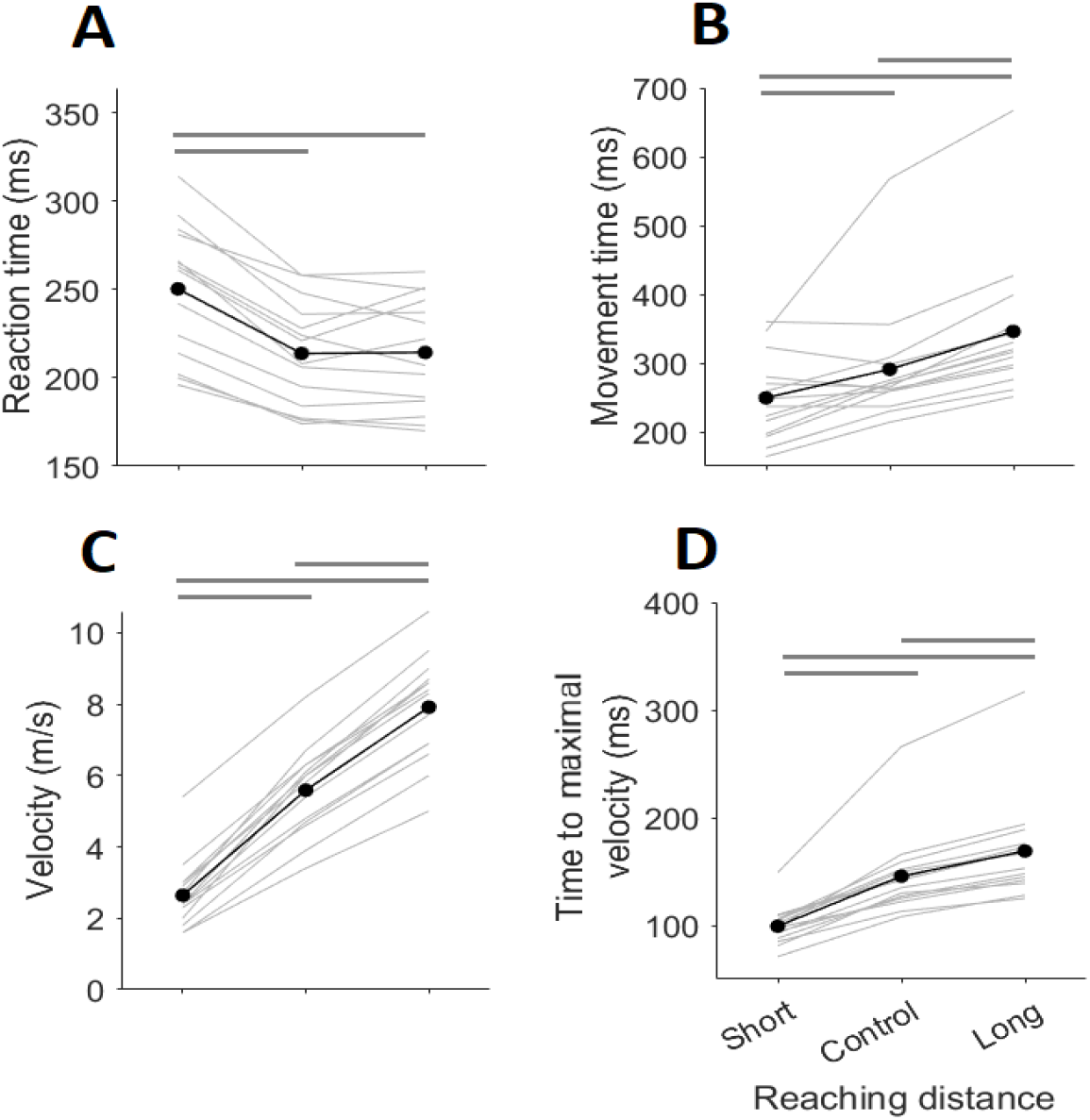
Condition-dependent variations of the reaction time (A), maximal velocity (B), movement time (C), time to the maximal velocity (D) and maximal velocity index within the entire movement (E) for the first experiment. On each plot, each thin light-grey line represents one of the 14 subjects who completed the first experiment, whereas the thick black dotted line represents the average across subjects. The horizontal thick dark-grey lines on top of the subjects represent the between-condition statistically significant (*p*<0.05) differences.

We and others have previously showed that larger express visuomotor responses are commonly associated with earlier RTs (Pruszynki et al., 2010; Gu et al., 2016; Contemori et al., 2021a). Here, we found that the task conditions leading to the latest (or earliest) RTs also led to the smallest (or largest) express visuomotor responses (see results). Critically, this calls into question whether the task-dependent modulation of express visuomotor response magnitude simply reflects the task-dependent generation of earlier or later reaches. To resolve this, we re-ran the detrended-integrated signal analysis on trial subsets with overlapping RTs (Dash et al., 2018; Kozak et al., 2019). We reasoned that if the express visuomotor response magnitude reflected task-dependent modulations of the express circuit, then similar between-condition contrasts should be observed in both original and RT-matched data sets. Note that, for each experiment, this additional analysis was run solely for the participants who exhibited express motor responses across all the three task conditions. For each of these subjects, we first pooled all the correct trials and generated the full list of RT values. We then ran an RT-match procedure by verifying the presence of at least one trial per condition for each RT±2ms value of the full data sample. Note that this approach was chosen to be conservative on the number of non-matching RTs to discard, which would otherwise increase by searching for perfect RT-match between conditions. Critically, this procedure generated three condition-specific data sets having similar RT-based distributions but different numbers of trials across conditions. To create compatible data sets, we binned the RT-matched trials every 20ms from the smallest RT value. Each bin of trials was then resampled with replacement 100 times by using a bootstrapping approach, and we selected an equal amount trials per condition based on the lowest number of trials across conditions for that bin in the original data set. In the end, we re-ran the detrended-integrated signal analysis methods on the RT-matched data sets.

#### Statistical analysis

Repeated measures ANOVA analyses with Bonferroni correction were conducted in SPSS (IBMSPSS Statistics for Windows, version 25, SPSS Inc., Chicago, Ill., USA) as the normality of the distributions was verified by the Shapiro–Wilk test. ANOVA analyses were conducted with task condition (3 levels: first experiment control long-reach, short-reach; second experiment control, overshoot, undershoot) as within-participant factors. When the ANOVA revealed a significant main effect or interaction, we ran Bonferroni tests for post-hoc comparisons. For all tests, the statistical significance was designated at *p*<0.05.

## RESULTS

### Experiment 1

#### Kinematic results

In the first experiment, the participants had to reach visual targets that could appear at different rightward or leftward distances from their dominant hand (for details see Experiment 1 task design). They successfully achieved the task goal in more than 90% of the trials across the three experimental conditions.

Figure 2A shows exemplar correct hand-to-target trajectories of a participant who completed the first experiment. For this subject, the targets requiring short reaching distances resulted in longer RT relative to control and long-reach conditions (dashed vertical lines in Figure 2B). After its initiation, the movement evolved at faster and slower velocities than control for the long-reach and short-reach conditions, respectively (dotted vertical lines in Figure 2B). The task-dependent variation in maximal velocity did not fully compensate that in reaching distance thus leading to longer movement times to complete longer than shorter reaches (Figure 2B). Note, however, that the participants were not required to complete the movement within a specific time (see Materials and Methods). Also, besides the larger time costs to reach higher velocity peaks, the hand velocity trace displayed a bell-shaped curve peaking at around the movement half across all conditions (Figures 2C).

For the entire group, the ANOVA showed statistically significant hand-to-target distance main effects for RT (*F_2,12_*=54.5, *p*<0.001), movement time (*F_2,12_*=37.8, *p*<0.001), maximal hand velocity (*F_2,12_*=250.9, *p*<0.001), time to maximal hand velocity (*F_2,12_*=35.5, *p*<0.001) and point of the maximal hand velocity within the movement (*F_2,12_*=8.9, *p*=0.004).

The short-reach target condition led to significantly longer RT than the other conditions (Figure 3A) and involved significantly lower maximal hand velocities (Figure 3C) although the reaching movement was completed significantly earlier (Figure 3B). Peak hand velocity was also reached significantly earlier than for the control condition (Figure 3D). By contrast, the long-reach target condition led to the opposite results. When the peak-velocity event was indexed relative to the whole movement duration, however, statistically significant differences (*p*<0.05) were observed solely between the short-reach (45±8% of the entire duration) condition and the other conditions (control, 50±6%; long-reach, 49±7% of the entire duration).

Overall, these results indicate that the participants were biased by the hand-to-target distances such that they took more time to start moving toward targets appearing close to their hand. Once the movement started, the hand velocity was modulated according to the hand-to-target distance but the greater hand speeds for longer reaches were insufficient to complete the task within the same time across conditions. Nevertheless, the hand was always accelerated for approximately half the movement distance before being decelerated to stop at the target, resulting in similar movement profiles for all hand-to-target distances.

#### EMG results

Figure 4 shows EMG data recorded from the PMch of an exemplar participant who met the conditions for positive express visuomotor response detection (see Materials and Methods) across all conditions of the first experiment. In the raster plot of Figure 4A-F, express visuomotor responses appear as a vertical band of either muscle activations (left targets) or inhibitions (right targets) at ~100 ms after the target presentation time. For this subject, the vertical band of EMG responses within the express time-window (i.e. 70-120 ms after the target onset; see Materials and Methods) became more evident as the hand-to-target distance increased (Figure 4A vs C vs E and B vs D vs F). Indeed, the number of trials with an express visuomotor response initiation increased by increasing the hand-to-target distance (see the red scatters and bars in Figures 4A-F). Specifically, the prevalence of express visuomotor response was ~60%, ~70% and ~80% for the short-reach, control, and long-reach conditions respectively. In addition to condition-dependent variation in express visuomotor response prevalence, we observed between-condition differences in express visuomotor response magnitude. Specifically, the average EMG signal enclosed in the express time-window (grey patch in Figure 4G) was smaller for the short-reach condition than the other conditions. By contrast, the earliest stimulus-driven EMG response was initiated ~90-100 ms after the stimulus onset across all conditions (inset plot in Figure 4G).

**Figure 4:**
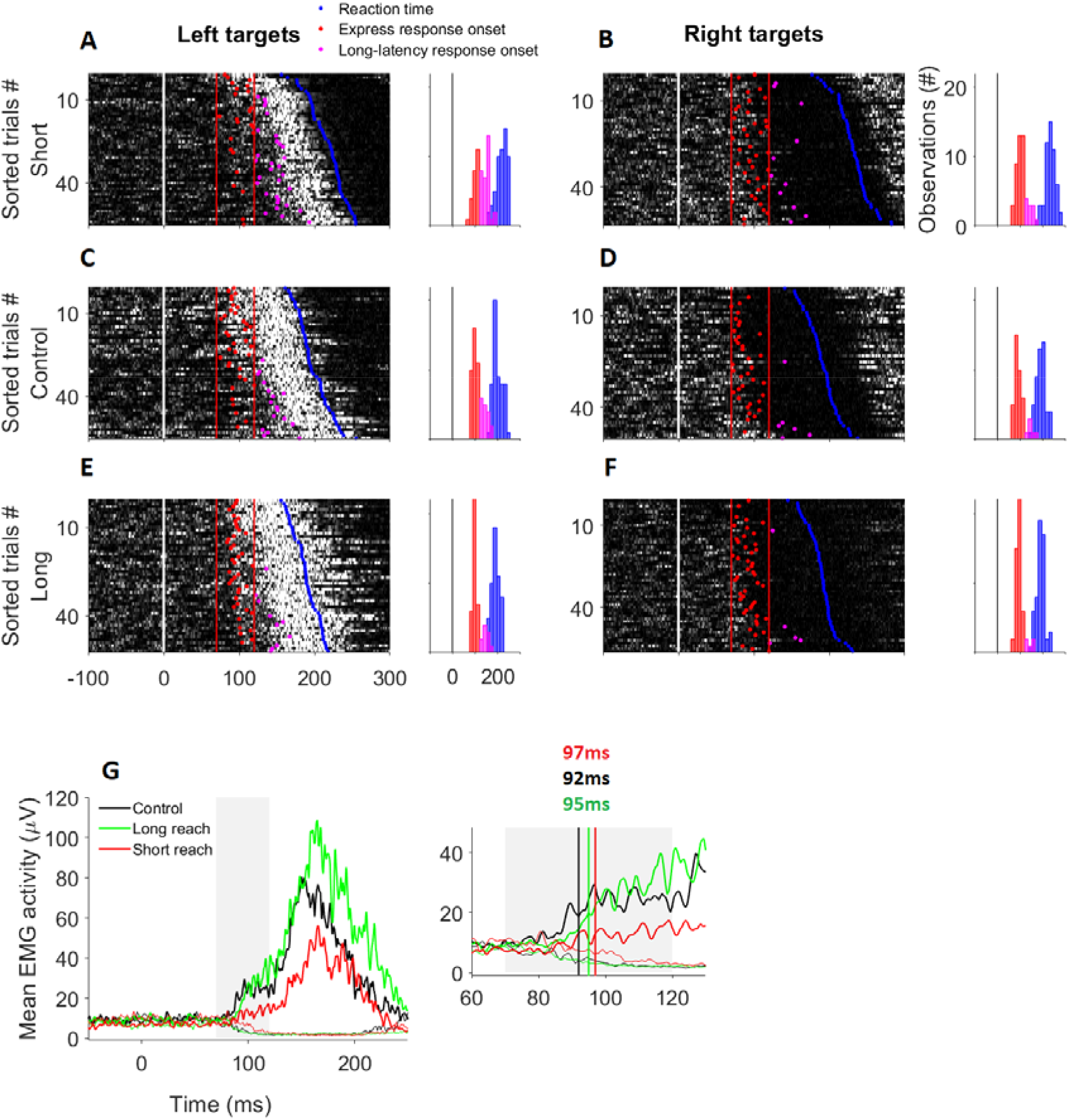
Surface EMG activity of the PMch muscle during the leftward and rightward movements executed toward the target requiring short (~3 cm; panels A and B), control (~8cm; panels C and D) and long (~13 cm; panels E and F) reaching distances of an exemplar participant who completed the first experiment and exhibited an express response in each of the three different hand-to-target distance conditions (see Materials and Methods). Each raster plot in panels A-F shows the rectified EMG activity from individual trials sorted by reaction time (brighter white colours indicate greater EMG activity). The white vertical line at 0ms indicates the target presentation time, the red vertical lines at 70 ms and 120 ms indicate the limits of the express visuomotor response time-window (see Materials and Methods), and the blue scatters indicate the reaction time. The express and long latency muscle response initiation times are represented with red and magenta scatters, respectively. In the inset left plots in panels A-F are shown the distribution of express (red histograms) and long-latency (magenta histograms) muscle response onset times as is that of reaction time (blue histograms).The task dependent average EMG activity computed across all trials is shown in panel G (thick lines = left target reaches; thin lines =right target reaches), as is a zoomed view of the mean EMG signal enclosed in the express time window (grey path at 70-120 ms). The vertical lines in the inset plot of panel G highlight the onset time (averaged across right and left target-directed express trials; see Materials and Methods) of express visuomotor response in the three task conditions.

For the first experiment, a total of ten participants (i.e. ~71% of the sample) exhibited express visuomotor responses on the PMch in all three conditions. For these subjects, we observed consistent condition-dependent variation of the express visuomotor response metrics. Indeed, the ANOVA showed statistically significant hand-to-target distance main effects for express visuomotor response prevalence (*F_2,8_*=57, *p*<0.001). The post-hoc analysis revealed that the prevalence of express visuomotor responses was significantly lower for the short reach than the other task conditions, and significantly higher than control for the long-reach condition (Figure 5A).

**Figure 5:**
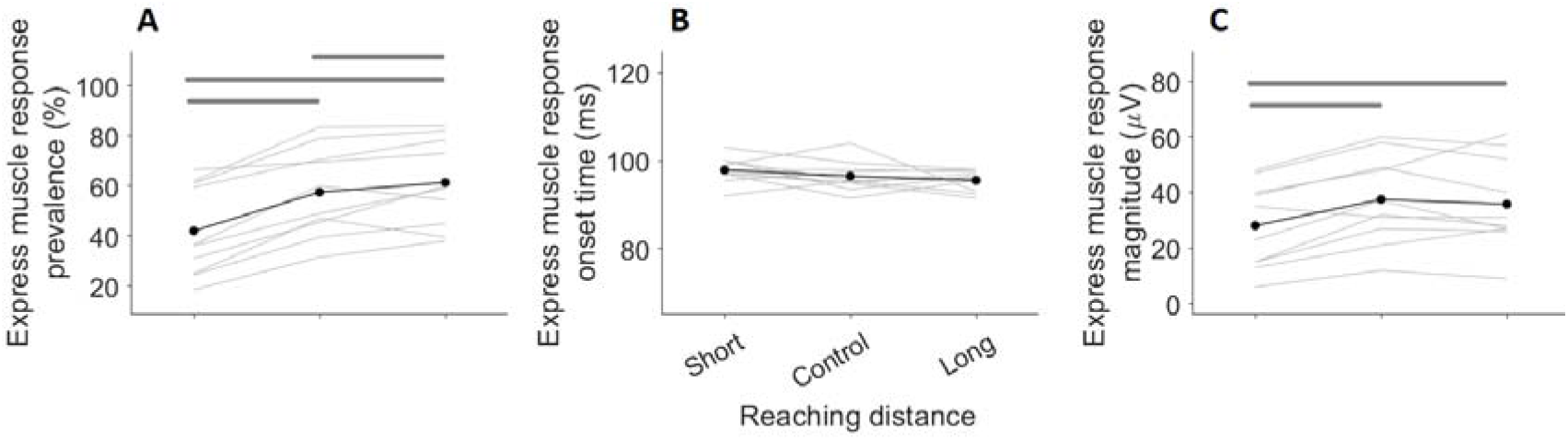
Condition-dependent variations of the prevalence (A), onset time (B) and magnitude (C) of express visuomotor responses. On each plot, each light-grey line represents one of the twelve subjects who exhibited the express response behaviour across all the three conditions of the first experiment, whereas the black dots represent the average across subjects. The horizontal dark-grey lines on top of the subjects represent the between-condition statistically significant (*p*<0.05) differences.

Although the express visuomotor response onset time tended to decrease with the hand-to-target-distance (Figure 5B), we did not find statistically significant contrast between the three conditions (*F_2,8_*=3.14, *p*=0.1).

The express visuomotor response magnitude was significantly modulated by the hand-to-target distance (*F_2,8_*=12.6, *p*<0.01). The post-hoc analysis revealed that the express visuomotor response magnitude was significantly smaller for the short reach than the other conditions (Figure 5C). Similar results were found also for the RT-matched data sets (short reach, ~28 μV; control and long reach, ~34 μV; target-distance main effect, *F_2,6_*=8.5, *p*<0.05; short vs control and short vs long, *p*<0.05; control vs long, *p*=0.08). Note that two participants were excluded from this analysis because the RT-match procedure (see Materials and Methods) discarded too many trials (~80%). This indicates that the express visuomotor response magnitude was modulated from trial to trial by the hand-to-target distance, independently of the time at which the volitional movement was initiated. It is also worth noting that these results are unlikely to reflect fixation-dependent differences in target perception, as target eccentricity was kept equal across the hemi visual fields for all task conditions (see Materials and Methods).

These results indicate that the express visuomotor response was modulated by the visuospatial task metrics such that targets reachable via small (or large) hand displacements inhibited (or facilitated) the generation of frequent and robust muscle responses within the express time limits. Note, however, that the earliest stimulus-driven muscle response was observed at similar times across conditions.

### Experiment 2

#### Kinematic results

The second experiment tested whether the context-dependent results of the first experiment reflected encoding of the veridical hand-to-target reaching metrics or preparation of movements of different amplitudes regardless of the real hand-to-target distance. Specifically, the participants were required to overshoot, undershoot, or stop at the target as a function of explicit trial-based instructions (for details see Experiment 2 task design). They successfully achieved the task goal in more than 90% of the trials across the three experimental conditions. For an exemplar subject, the requirement to stop at the actual target location (i.e. control condition) resulted in earlier RTs relative to the other tasks (dashed vertical lines in Figure 6B). Higher velocity and longer movement time were observed for the overshoot than control conditions, whereas the undershoot task led to the opposite results (Figure 6B). Also, the hand was accelerated for longer in conditions with higher peak velocities (Figure 6B). The movement, however, evolved similarly across conditions with a single acceleration phase terminating within the first half of the movement (Figures 6C).

**Figure 6:**
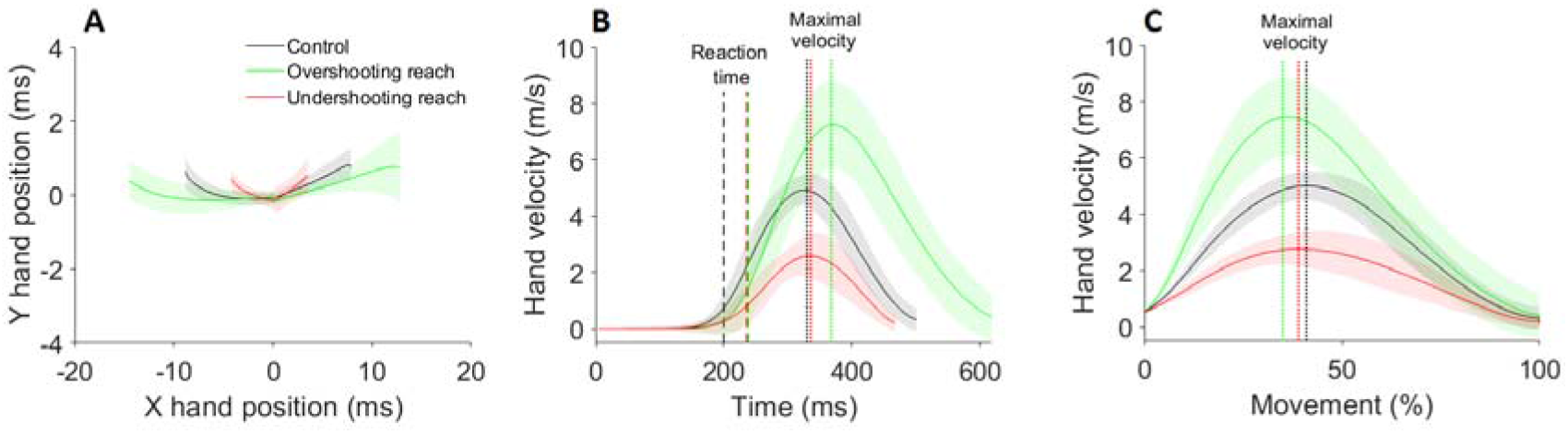
Task-depended hand trajectories (A), velocity traces (B) and time-normalized hand velocity traces showing the point at which the hand reached the peak velocity within the entire movement (C) of an exemplar participant for the second experiment (same format as Figure 2).

For the entire group, the ANOVA showed statistically significant task-condition main effects for RT (*F_2,10_*=14.9, *p*<0.001), movement time (*F_2,10_*=61.7, *p*<0.001), maximal hand velocity (*F*_2,10_=190, *p*<0.001), time to maximal hand velocity (*F_2,10_*=105.2, *p*<0.001) and point of the maximal hand velocity within the movement (*F_2,10_*=13.9, *p*=0.001). The movement started significantly earlier in the control condition than the other conditions (Figure 7A) and ended significantly earlier and later than control for the undershoot and overshoot conditions, respectively (Figure 7B). The hand moved significantly slower than control for the undershoot and significantly faster than control for the overshoot conditions (Figure 7C). Peak velocity occurred significantly earlier than control for the undershoot and significantly later than control for the overshoot conditions (Figure 7D). The peak velocity occurred at around half of the movement distance for the control (50±7% of the entire duration) and overshoot conditions (49±8% of the entire duration), but significantly earlier (*p*<0.05; 46±6% of the entire duration) than this for the undershoot conditions.

**Figure 7:**
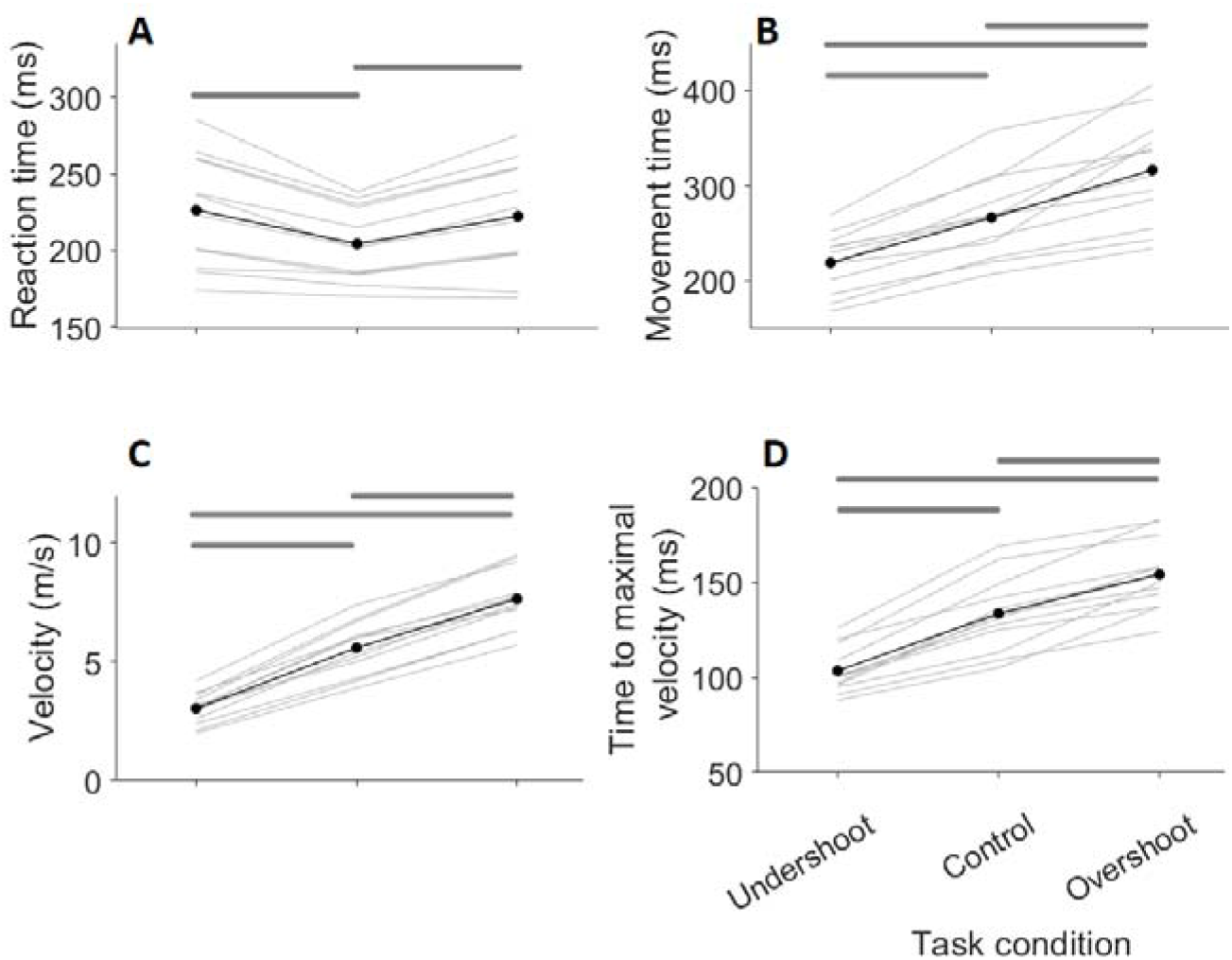
Task-dependent variations of the reaction time (A), maximal velocity (B), movement time (C), time to the maximal velocity (D) and maximal velocity index within the entire movement (E) for the second experiment (same format as figure 3).

In all, these data show that the requirement to not stop at the actual target location increased the RT by a similar amount for both the overshooting and undershooting tasks. Once the movement started, however, it evolved similarly across conditions besides expectable task-dependent variations in movement velocity and time. These results might indicate that the movement endpoint was encoded during the premotor planning process, which was complicated by mismatching the target and task-goal metrics (e.g. stop before\after the target).

#### EMG results

Figure 8 shows exemplar EMG data recorded from the PMch of a participant who exhibited express visuomotor responses across all the three conditions of the second experiment (same subject as Figure 4). For this subject, the number of trials with an express visuomotor response was larger in the control than the other task conditions (see the red scatters and the bars in Figures 4A-F). Indeed, the prevalence of express visuomotor response was ~72% for the control condition and ~45% for the overshoot and undershoot conditions. Consistently, the prevalence of long-latency muscle responses was smaller for the control than the other conditions (see the magenta scatters and bars in Figures 8A-F).

**Figure 8:**
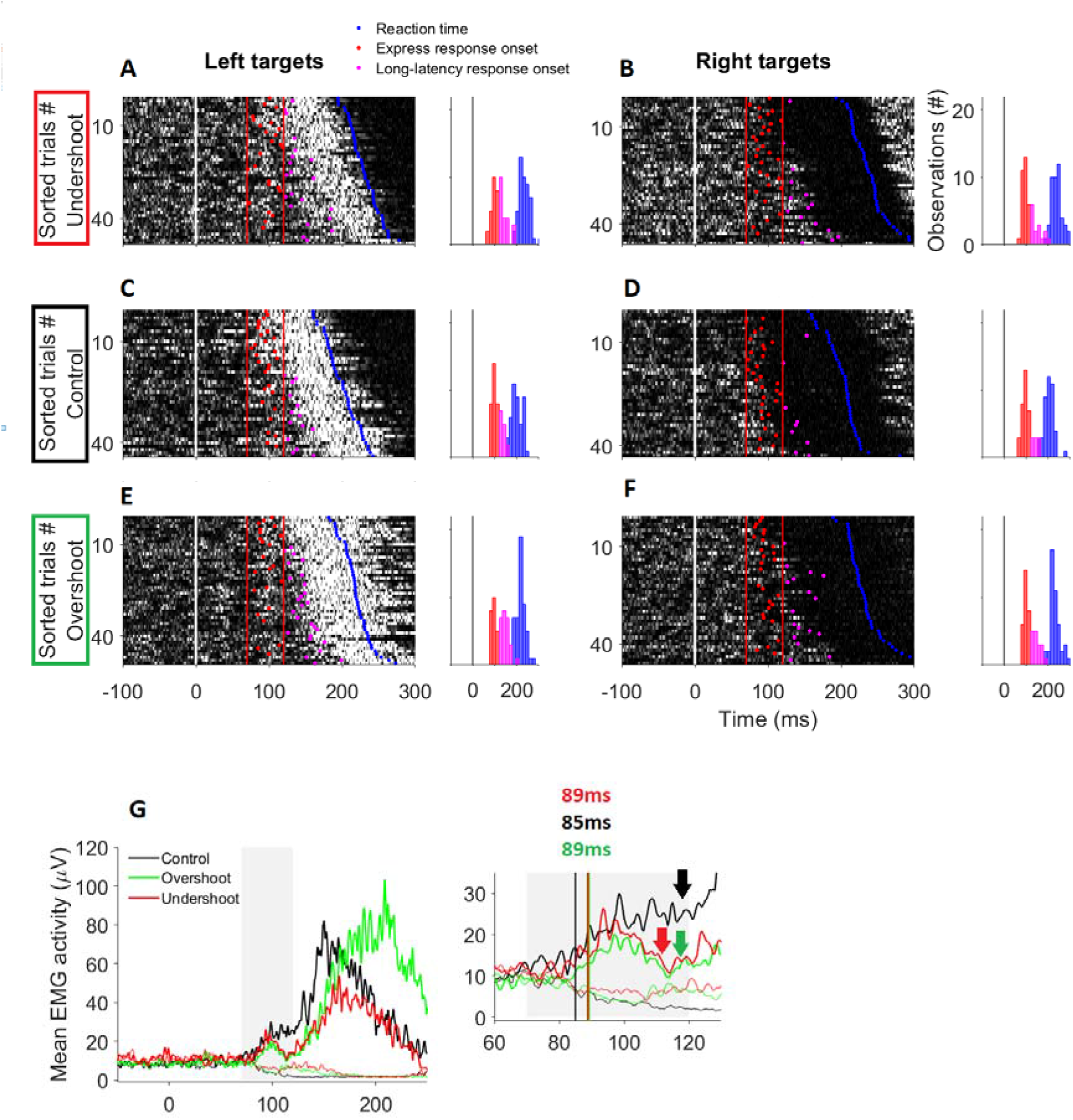
Surface EMG activity of the PMch muscle during the undershooting (panels A and B), control (i.e. stop at the target; panels C and D) and overshooting (panels E and F) visuospatial tasks of an exemplar participant who completed the second experiment. Rasters of rectified EMG activity from individual trials are shown in panels A-F, as are the distributions of express and long-latency visuomotor response onset times and reaction time relative the target presentation time. The task-dependent average EMG activity computed across all trials is shown in panel G, as are a zoomed view of the mean EMG signal enclosed in the express time window and the onset time of express visuomotor response in the three task conditions. In the inset plot of panel G, the arrows highlight the transition of the EMG signal across the express and long-latency epochs. Panels A-G have the same format as figure 4.

The express visuomotor response magnitude was larger and started earlier for the control than the overshoot and undershoot target conditions (Figure 8G). Notably, the long-latency EMG signal was superimposed on the express EMG signal without a clear separation between the two phases for the control condition (see the black arrow in the inset plot of Figure 8G), suggesting close temporal summation of congruent express and long-latency signals. By contrast, for both the overshoot and undershoot conditions the express EMG signal returned close to the background level prior to initiation of the long-latency response (see the green and red arrows in the inset plot of Figure 8G).

For the second experiment, ten out of the twelve participants (i.e. ~83% of the sample) met the conditions for positive express visuomotor responses detection on the PMch across the three conditions. For these subjects, the ANOVA showed statistically significant task-condition main effects for express muscle response prevalence (*F_2,8_*=8.2, *p*<0.05), which was significantly larger in the control than the other task conditions (Figure 9A). The express visuomotor response tended to initiate earlier for the control than the other conditions (Figure 9B), but this between-condition contrast was not statistically significant (*F_2,8_*=3.4, *p*=0.09).

**Figure 9:**
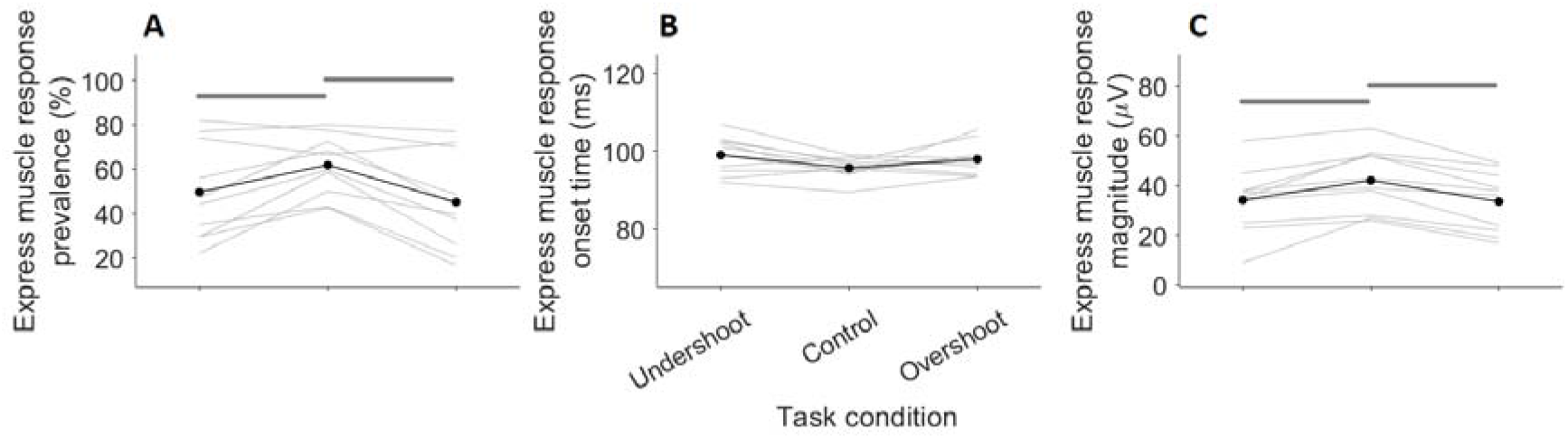
Task-dependent variations of the prevalence (A), onset time (B) and magnitude (C) of express visuomotor responses (same format as figure 5).

The express visuomotor response magnitude was significantly modulated by the task conditions (*F_2,8_*=27, *p*<0.001) as it was significantly smaller than control for both the overshoot and undershoot task conditions (Figure 9C). Notably, similar statistical results were found also for the RT-matched data sets (control, ~38 μV; overshoot and undershoot, ~32 μV; task-condition main effect, *F_2,6_*=11.7, *p*<0.01; control vs undershoot and control vs overshoot, *p*<0.05; undershoot vs overshoot, *p*=0.9). This was consistent with the first experiment findings and suggests the express response magnitude was systematically modulated by the trial-based context, irrespective of the RT. Again, two participants were excluded from this analysis because of an excessive (>80%) trial discarding rate during the RT-match procedure (see Materials and Methods).

These results indicate that matching the real target with the task-goal endpoint facilitates express transformation of visual inputs into appropriate motor outputs, relative to when the movement had to stop at a non-veridical target location. When express responses occurred, however, their processing times were similar for real (veridical) and abstract (non-veridical) target locations.

## DISCUSSION

This study showed that express visuomotor responses reflect encoding of both the physical hand-to-target distance and intended movement amplitude during a visuospatial task. This suggests that the express visuomotor outputs can be strategically exploited by the cerebral cortex to facilitate rapid and appropriate response initiation during target-directed actions. A schematic representation of a possible circuit organisation is outlined in Figure 10.

**Figure 10:**
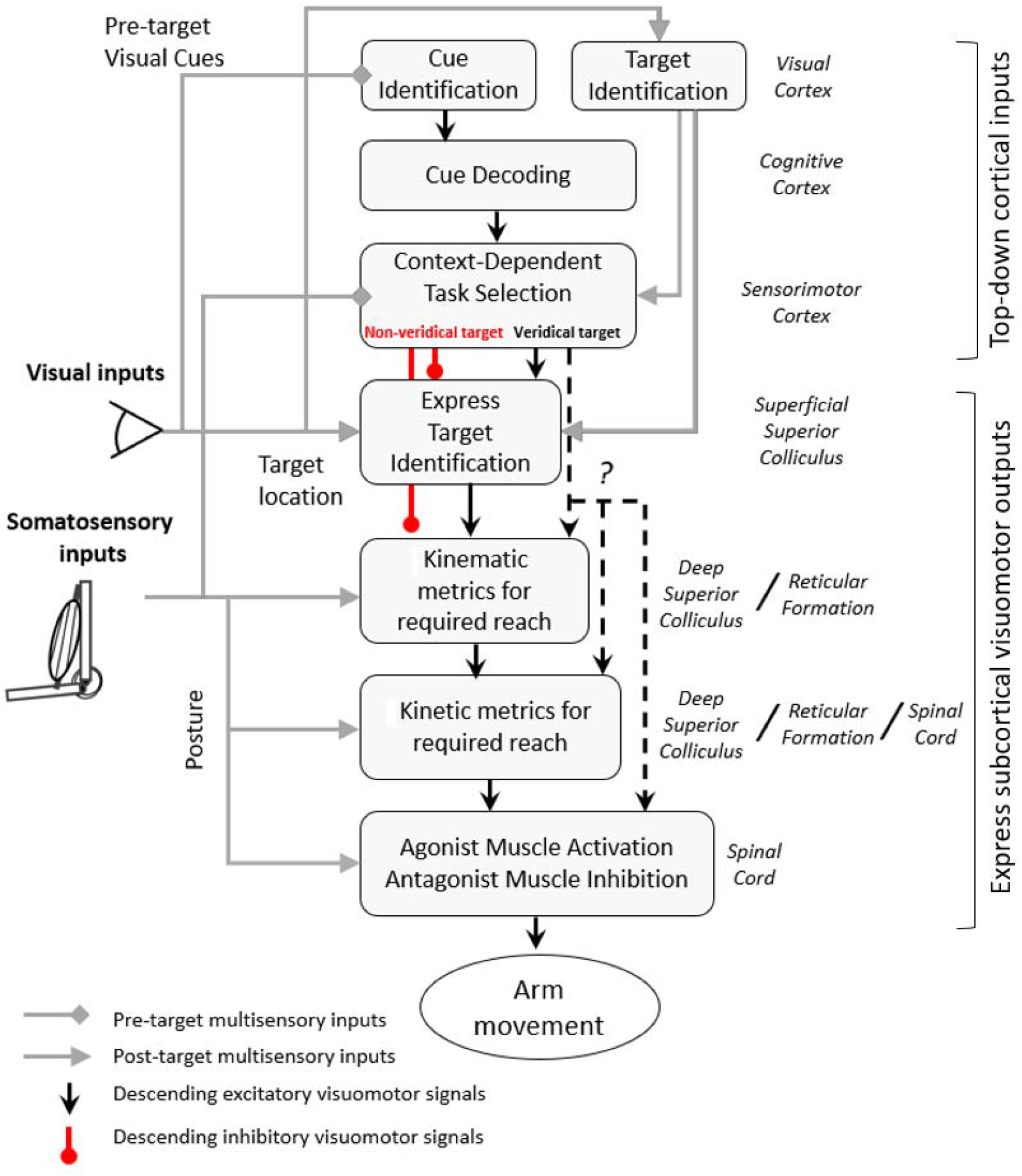
Proposed cortico-subcortical coordination for visuomotor response generation in which cerebral cortex areas inhibit or facilitate the express tecto-reticulo-spinal system according to whether the task involves veridical or non-veridical targets to reach. The reach-goal location resulting from integration of physical target location and cued instructions is converted into the extent of the reach based on multisensory signalling of the current arm position (kinematics), the required joint torques (kinetics) and muscle activation via spinal interneurons and motoneurons. Dashed black lines with ‘?’ mark denote uncertainty about whether the kinematic-kinetic metrics for the later (voluntary) reach component is computed by a transcortical or subcortical network.

### Mechanisms for express visuomotor responses during reaches to veridical targets

The first experiment showed that the physical hand-to-target distance modulated express visuomotor transformations ascribable to the subcortical tecto-reticulo-spinal circuit. Notably, the involvement of the superior colliculus in the generation of express visuomotor responses (Corneil et al., 2004, 2008; Pruszynski et al., 2010) is consistent with its capability to encode the location of either target or distractor stimuli within ~40–70ms (Boehnke and Munoz, 2008). Further, the surface layer of this midbrain structure is organized in retinotopic map, whereas its deeper layers are organized in somatotopic maps (for review see Basso and May, 2017; Boehnke and Munoz, 2008). Therefore, the superior colliculus could integrate visual and somatosensory information to compute the direction and distance of a target-directed action. Downstream from the superior colliculus, the reticular formation also receives inputs from somatosensory afferents (Leiras et al., 2010). Thus, a tecto-reticulo-spinal pathway has the required sophistication to compute and generate appropriate express visuomotor responses during veridical target-directed actions of different amplitudes (Figure 10).

We recently documented circumstances of express visuomotor response modulation secondary to explicit cues (Contemori et al., 2021a, 2021b, 2022). Pre-stimulus information might facilitate processing of expected stimuli at the superficial superior colliculus (for review see Corneil and Munoz, 2014) and/or initiation of expected responses along the express pathway (Basso and Wurtz, 1998; Cisek and Kalaska, 2005; Contemori et al., 2022). Here, the eccentricity (hence saliency) of opposite targets was constant but local thresholds for responding could be modulated asymmetrically by corticotectal projections (Boehnke and Munoz, 2008). Alternatively, or additionally, the advance information about the required distance for each possible target could bias the tecto-reticulo to generate larger express signals for targets requiring larger movements. These, in turn, would be more likely to cross the spike threshold for neurons along the express pathway, as consistent with the increased express visuomotor response magnitude and prevalence associated with increased physical hand-to-target distance. Such a mechanism would also be expected to result in slightly shorter latencies for the larger responses, but the differences would be difficult to detect from these EMG signals.

Previous work suggests that the express responses contribute to the volitional visuomotor behaviour because larger express outputs were associated with earlier RTs (Pruszynki et al., 2010; Gu et al., 2016; Contemori et al., 2021a). Mechanically, RT detection depends on muscle force accelerating the arm to the threshold velocity. Notably, the muscle force will rise more rapidly if the same motor units receive close temporospatial summation of express and long-latency motor signals, which would enhance intramuscular calcium release and diffusion (i.e. the *catch* property of muscle; see for review Tsianos and Loeb, 2017). In our veridical reaching conditions, the express visuomotor responses prevalence and amplitude were associated with the amplitude of the long-latency EMG signal (>120ms; Fig. 4G) that reflected the required reaching length. As expected, summation of larger express and long-latency muscle recruitment generated earlier RTs and higher peak velocities. It is also possible that the weaker express visuomotor responses associated with shorter reaches reflect, at least partially (see Wong et al., 2017), the higher complexity (i.e. RT delay) inherent in planning short movements that offer less time for online correction.

### Mechanisms for express visuomotor responses during reaches to non-veridical targets

In the second experiment, the deliberate intent to reach toward non-veridical targets in the overshoot and undershoot tasks led to reduced prevalence and magnitude of express visuomotor responses relative to control. Why should the express response be inhibited in circumstances requiring non-veridical responses?

The fact that express visuomotor responses rigidly encode the visual stimulus location (Wood et al., 2015; Gu et al., 2016; Atsma et al., 2018) is consistent with their proposed subcortical origin (Pruszynki et al., 2010). Nevertheless, this also suggests that the express motor output might be counterproductive when the task requires non-veridical responses. For instance, Gu et al., (2016) showed that large pro-target express responses delayed the initiation of correct anti-target reaches, plausibly because of larger time costs to override the express pro-target muscle forces (Gu et al., 2016). The express system, however, appears to be flexible to contextual task rules when these are predictable. For instance, Wood and colleagues (2015) recorded express target-directed muscle responses in delayed-reach task trials that were randomly intermingled with no-delay trials (i.e. task condition unpredictability). By contrast, express visuomotor responses were obliterated when only delayed-reach trials were presented within a block (Pruszynski et al., 2010). Notably, these results are consistent with more recent evidence of cortico-subcortical modulation mechanisms for express visuomotor responses as a function of explicit cues (Contemori et al., 2021a, 2021b, 2022). Considering that express visuomotor responses aid rapid movement initiation (Pruszynki et al., 2010; Gu et al., 2016; Contemori et al., 2021a), their inhibition could reflect strategic cortico-subcortical inhibition in contexts requiring longer RTs, such as for reaching toward non-veridical targets (Figure 10).

The overshoot/undershoot tasks resulted in longer RTs than control, which could reflect increased complexity to plan a non-veridical response trajectory (Wong et al., 2016). Cortical planning of appropriate responses for achieving complex task goals (e.g. reaching an abstract target) can modulate the networks downstream from the cerebral cortices (Selen et al., 2012; see for review Kurtzer, 2015). Critically, these include the reticulo-spinal circuits that are proposed to process the superior colliculus signals to generate express visuomotor responses (Corneil et al., 2004, 2008; Pruszynski et al., 2010; Figure 10).

The mammalian reticular formation is involved in the control of static posture (Sherrington, 1898; Rhines and Magoun, 1946; Magoun and Rhines, 1946). More recent neurophysiological and behavioural work also suggest that this brainstem structures contributes to volitional upper-limb movements (Alstermark and Isa, 2012; Contemori et al., 2021c) and reflexive responses to mechanical perturbations of static upper-limb postures (Kurtzer, 2015). Notably, the reticular formation receives descending signals from both the superior colliculus (Boehnke and Munoz, 2008) and cortical brain areas (Keizer and Kuypers, 1984, 1989; Fregosi et al., 2017; Darling et al., 2018; Fisher at al. 2021). The reticular formation, therefore, appears to be well-placed to integrate descending collicular signals encoding the physical stimulus location (Everling et al., 1999; McPeek and Keller, 2002) with cortical premotor signals affording task-related rules (e.g. to overshoot\undershoot the target). In this circumstance, express stimulus-driven motor signals may be inhibited (or even obliterated) to delay the RT when there is uncertainty about the reach goal, such as for our non-veridical reaching tasks (Figure 10).

### A common subcortical network for rapid initiation and online control of reaching?

Both the mammalian superior colliculus (Werner, 1993) and reticular formation (Buford and Davidson, 2004; Schepens and Drew, 2004) are active before and during upper limb reaching movements. This suggests that the tecto-reticulo circuit might contribute to both the initiation (Pruszynki et al., 2010; Gu et al., 2016; Contemori et al., 2021a) and online control of reaching movements (Day and Lyon, 2000; Day and Brown, 2001). Notably, this is consistent with recent evidence of express visuomotor responses to correct the online movement trajectory in a jump-target task (Kozak et al., 2019). Furthermore, the size of muscle responses starting ~90-120 ms after visual perturbation of virtual hand position showed a non-linear scaling for perturbation amplitudes >2 cm (Cross et al., 2019). Here, we observed a similar non-linear scaling of the express visuomotor response magnitude although for hand-to-target distances >8 cm, probably because the participants responded from static rather than dynamic (Cross et al., 2019) postures.

Initiation and online control of real-world visuospatial actions rely on visuomotor circuits that must integrate multisensory information about the body and target positions, which is inherently variable and noisy. The current data is consistent with previous evidence suggesting that the putative subcortical express circuits can be primed to generate flexible context-dependent motor outputs that support the accomplishment of visuospatial tasks from both static (Kurtzer, 2015; Weiler et al., 2019; Contemori et al., 2022) and dynamic postures (Cross et al., 2019; Kozak et al., 2019; Weiler et al., 2021).

### Conclusions

This study shows that express visuomotor responses can be flexibly modulated to achieve visuospatial task-goals. The data are consistent with the idea of a subcortical visuomotor pathway whose motor output is strategically exploited by the cerebral cortex to facilitate rapid initiation of veridical target-directed reaches. It remains to be determined whether the longer latency, presumably cortically-driven, motor responses rely on some or all of the same subcortical circuits to convert reaching targets in extra-personal space coordinates into patterns of muscle activity that will achieve the desired limb movements. Overall, our findings emphasise the need for consideration of subcortical sensorimotor circuits in theories of human motor control and behaviour.

## Acknowledgements

This work was supported by operating grants from the Australian Research Council (DP170101500) awarded to T.J. Carroll, B.D. Corneil, G.E. Loeb and G. Wallis.

